# Loss of intermicrovillar adhesion impairs basolateral junctional complexes in transporting epithelia

**DOI:** 10.1101/2024.03.19.585733

**Authors:** Caroline S. Cencer, Kianna L. Robinson, Matthew J. Tyska

**Author notes:** To whom correspondence should be addressed: Matthew J. Tyska, Ph.D., Department of Cell and Developmental Biology Vanderbilt University School of Medicine, T-2212 Medical Center North, 465 21st Avenue South Nashville, TN 37240-7935. Abbreviations: TJ, AJ, IMAC, CL4, DPC, KO, TEER.

## Abstract

Transporting epithelial cells in the gut and kidney rely on protocadherin-based apical adhesion complexes to organize microvilli that extend into the luminal space. In these systems, CDHR2 and CDHR5 localize to the distal ends of microvilli, where they form an intermicrovillar adhesion complex (IMAC) that links the tips of these structures, promotes the formation of a well-ordered array of protrusions, and in turn maximizes apical membrane surface area. Recently, we discovered that IMACs can also form between microvilli that extend from neighboring cells, across cell-cell junctions. As an additional point of physical contact between cells, transjunctional IMACs are well positioned to impact the integrity of canonical tight and adherens junctions that form more basolaterally. Here, we sought to test this idea using cell culture and mouse models that lacked CDHR2 expression and were unable to form IMACs. CDHR2 knockout perturbed cell and junction morphology, led to loss of key components from tight and adherens junctions, and impaired barrier function and wound healing. These results indicate that, in addition to organizing apical microvilli, IMACs provide a layer of cell-cell contact that functions in parallel with canonical tight and adherens junctions to support the physiological functions of transporting epithelia.

## INTRODUCTION

Cell junctions are essential for the integrity of epithelial tissues, forming physical connections between neighboring cells that provide structural support and control paracellular permeability (Horowitz et al., 2023). By partitioning apical and basolateral components in the plane of the plasma membrane, junctional complexes also play an essential role in the acquisition and maintenance of cell polarity, which in turn defines a functional axis at the cell and tissue scales (Buckley and St Johnston, 2022). Transporting epithelia found in the small intestine and kidney tubules are two examples of tissues that rely heavily on junctional complex formation (Garcia-Castillo et al., 2017). In this context, cell junctions are organized into stacked layers consisting of (from apical to basal) the tight junction (TJ), the adherens junction (AJ), desmosomes, gap junctions, and hemidesmosomes (Garcia et al., 2018). Loss of key components from these distinct layers results in broad ranging defects in cell and tissue architecture and function (Buckley and St Johnston, 2022; Horowitz et al., 2023).

At the level of the TJ, transmembrane proteins including junctional adhesion molecule (JAM), occludins, and claudins create adhesive strand-like structures that interact with TJ components on the opposing surface of a neighboring cell (Van Itallie and Anderson, 2014). Inside the cell, zonula occludens (ZO-1,-2,-3) act as scaffolding proteins to couple TJ adhesion molecules to F-actin and non-muscle myosin 2, which provide peripheral support and generate mechanical tension (Horowitz et al., 2023). TJs are responsible for selectively controlling solute permeability in different tissue contexts, with specific complements of claudin isoforms creating distinct permeability profiles based on particle size and charge (Tsukita et al., 2019).

Immediately basal to TJs are AJs, which are defined by high levels of E-cadherin, a single-spanning transmembrane protein that drives strong adhesion between neighboring cells (Troyanovsky, 2023). Knockout (KO) of E-cadherin in mouse intestinal tissues demonstrated clear roles in intestinal morphogenesis and barrier function with blunted villi and increased TJ permeability (Bondow et al., 2012). β-catenin is a core cytoplasmic component of the adherens junction, which binds to E-cadherin and through an interaction with α-catenin, forms a physical link to the underlying actin cytoskeleton (Tian et al., 2011; Valenta et al., 2012). Loss of E-cadherin releases β-catenin from adherens junctions, leading to activation of the Wnt-signaling pathway, with downstream loss of the epithelial phenotype and enhanced cell migration, as observed in a broad range of cancers (Tian et al., 2011; van der Wal and van Amerongen, 2020; Yap, 1998). While the factors alluded to above are well studied in the context of junctional biology, new roles and localizations for these components are still being discovered. For example, polarity proteins including CRB3A, PAR6β, and aPKC, were recently localized along the length of apical microvilli, outside of the tight junction (Mangeol et al., 2022). Furthermore, Nectin-3, an adhesion protein classically categorized as part of the AJ, has now been identified in microvilli in human colonic tissue and cultured cells (Childress et al., 2023; Mangeol et al., 2024).

Outside of the large collection of adhesion molecules that accumulate in junctional complexes, other adhesive factors play key roles in shaping epithelial cells. Transporting cell types, like those found in the kidney and gut, leverage intermicrovillar adhesion complexes (IMACs) to drive apical brush border assembly (Cencer et al., 2023; Crawley et al., 2014; Pinette et al., 2019). Here, protocadherins CDHR2 and CDHR5 target to the distal tips of protrusions, where they form heterophilic contacts that define the spacing between microvilli and maximize the number of protrusions that extend from the surface (Crawley et al., 2014; Pinette et al., 2019). Unexpectedly, recent studies also established that CDHR2 and CDHR5 form complexes between the tips of microvilli that extend from neighboring cells, forming ‘transjunctional’ IMACs that are extremely long-lived. These transjunctional complexes provide an anchoring mechanism for nascent microvilli and promote the long-term accumulation of protrusions on the apical surface (Cencer et al., 2023).

By providing an additional mode of physical contact between adjacent epithelial cells, transjunctional IMACs are well positioned to support the integrity of the more basolateral, canonical junctional complexes (e.g. TJs and AJs). To test this hypothesis, we examined multiple CDHR2 loss-of-function models and explored the possibility of junctional phenotypes in these systems. Here, we report that epithelial cell culture and mouse models lacking CDHR2 exhibit prominent defects in cell junctions, including lower levels of mechanical tension and a loss of established junctional components. These perturbations are accompanied by increased permeability and a highly motile phenotype.

We propose that, under normal conditions, these defects are prevented by CDHR2-dependent formation of transjunctional IMACs. More generally, these data suggest that transjunctional IMACs form a previously unrecognized layer of the junctional complexes that promote cell-cell adhesion.

## RESULTS AND DISCUSSION

### Loss of CDHR2 disrupts epithelial cell morphology

To characterize the potential impact of CDHR2 loss-of-function on canonical junctional complexes, we revisited our previously described CDHR2 KO LLC-PK1-CL4 (CL4) cell line (Cencer et al., 2023) and also developed a new KO model based on the intestinal epithelial CACO-2_BBE_ cell line (**Figure S1A**). Our previous study revealed that CDHR2 KO CL4 cells exhibit a striking loss of microvillar clustering and reduced accumulation of protrusions at cell margins (Cencer et al., 2023), as expected given the central function of this protocadherin in the IMAC (Crawley et al., 2014; Pinette et al., 2019). Analysis of CDHR2 KO CACO-2_BBE_ cells revealed a similar phenotype, with a lack of microvilli clustering, even late in differentiation (see 12 days post confluency, DPC) (**Figure S1B**). As with CDHR2 KO CL4 cells, levels of CDHR5 were also decreased at 12 DPC (**Figure S1C-E**), a phenotype consistent with other IMAC loss-of-function models (Choi et al., 2020; Crawley et al., 2016; Li et al., 2017; Pinette et al., 2019; Weck et al., 2016).

Further characterization of our existing and newly developed CDHR2 KO cell lines revealed highly aberrant cell morphologies and junctional profiles. Confocal microscopy of CDHR2 KO CL4 cells stained for tight junction component ZO-1 revealed elongated cell shapes with larger overall areas when viewed *en face* (**Figures 1A-C and S2B-C)**. CDHR2 KO CACO-2_BBE_ cells also displayed deformed junctions with a striking ’ruffled’ appearance (**Figure 1D, Zoom)**. Straightness measurements revealed that junctions in this KO line were less linear, with a mean straightness of 0.70 ± 0.13 compared to control junctions of 0.93 ± 0.03 (1 being a perfectly straight line; **Figure 1E**). Previous work established that junctional straightness is linked to the enrichment of non-muscle myosin-2 (NM2) (Choi et al., 2016; Tokuda et al., 2014; Van Itallie et al., 2009) and presumably high levels of tension in the contractile ‘belt’ that encircles the cell at the level of junctional complexes. Interestingly, apical cell area (**Figure 1A,B**) is also restricted by junctional tension, as reduced tension causes cells to spread (Diz-Munoz et al., 2013; Sumi et al., 2018). With these points in mind, we sought to determine if the loss of CDHR2 affected junctional tension by staining for non-muscle myosin-2C (NM2C), the most abundant NM2 variant in mature enterocytes (Chinowsky et al., 2020; Ebrahim et al., 2013). Staining for NM2C in fully differentiated (21 DPC) CDHR2 KO CACO-2_BBE_ cells revealed significantly lower levels of NM2C signal (**Figure 1F-G**), suggesting that junctional tension is reduced in these cells. Together, these results indicate that CDHR2, which localizes primarily to the distal ends of brush border microvilli, has roles in maintaining normal epithelial cell and monolayer morphology.

**Figure 1.**
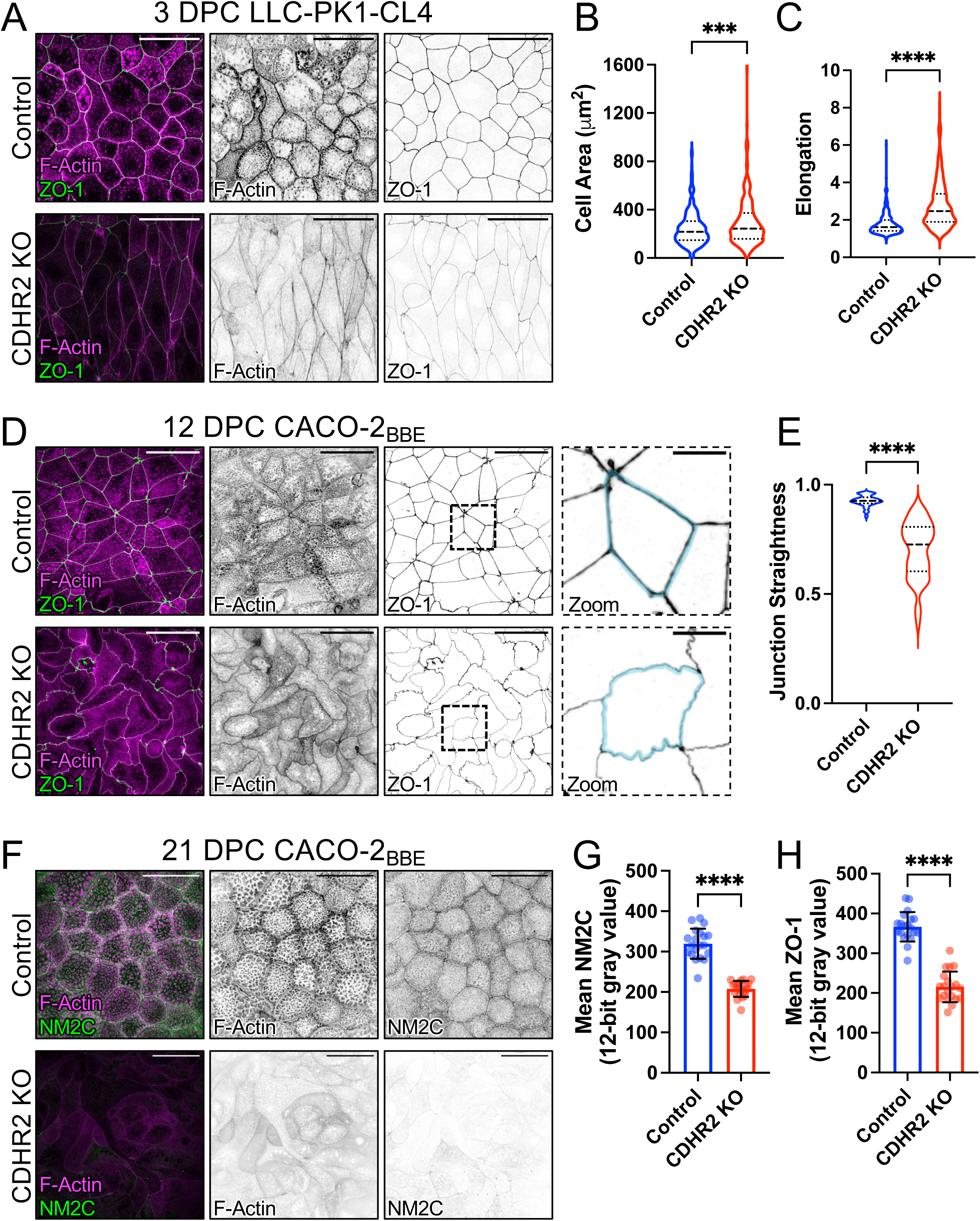
CDHR2 KO cells exhibit aberrant cell morphologies and decreased apical junction markers. (A) 3 DPC Control and CDHR2 KO CL4 cells stained for F-actin (magenta) and ZO-1 (green). (B) Cell area measured in n = 384 control cells and n = 259 KO cells. (C) Cell elongation measured as the ratio of max feret length to min feret length from the cells in (B). (D) 12 DPC Control and CDHR2 KO CACO-2_BBE_ cells stained for F-actin (magenta) and ZO-1 (green). Zooms show straight vs ruffled junctions in control and KO, respectively, taken from the dashed boxes. (E) Junctional straightness (see methods) from n = 62 control and KO cell junctional segments where 1.0 is a straight line. (F) 21 DPC Control and CDHR2 KO CACO-2_BBE_ cells stained for F-actin (magenta) and NM2C (green). (G) Mean NM2C and (H) ZO-1 intensities measured in CACO-2_BBE_ cells from n = 30 60X fields per condition. Unpaired t-tests; ***p = 0.003, ****p<0.005. Error bars represent mean ± SD. Scale bars: 40 µm (A,D,F), 10 µm (D, Zooms).

### Loss of CDHR2 perturbs the composition of canonical junctional complexes

The abnormal cell morphology and perturbations in the circumferential actomyosin belt induced by loss of CDHR2 (**Figure 1**) could be due to defects in the canonical junctional complexes that mediate cell-cell contact. Indeed, confocal imaging of ZO-1 staining, which we used to delineate cell boundaries in monolayers (**Figure 1A**), revealed reduced ZO-1 levels in both CDHR2 KO CL4 and CACO-2_BBE_ lines (**Figures 1H, S2C**), indicating defective TJ assembly in both models. Other basolateral proteins that directly contribute to cell-cell contacts, such as claudin-7 and epithelial cellular adhesion molecule (EpCAM), were also depleted in CDHR2 KO CACO-2_BBE_ cells (**Figure 2A-B, E-F**). As direct binding partners, claudin-7 and EpCAM contribute to TJ barrier maintenance through damage-induced cleavage of EpCAM and release of claudin-7 (Higashi et al., 2023; Ladwein et al., 2005; Wu et al., 2013). We also observed a significant loss of canonical AJ markers, including E-cadherin and β-catenin, when staining CDHR2 KO CACO-2_BBE_ cells (**Figure 2C-D, G-H**); these phenotypes were recapitulated in the CDHR2 KO CL4 model (**Figure 2I-K**). Importantly, ZO-1 and CDHR5 (heterophilic binding partner for CDHR2 in the IMAC) signals were partially rescued upon exogenous stable expression of CDHR2-HALO protein (**Figure S2A-D**), suggesting that these perturbations are due to the loss of CDHR2 in this KO line, rather than off-target effects.

**Figure 2.**
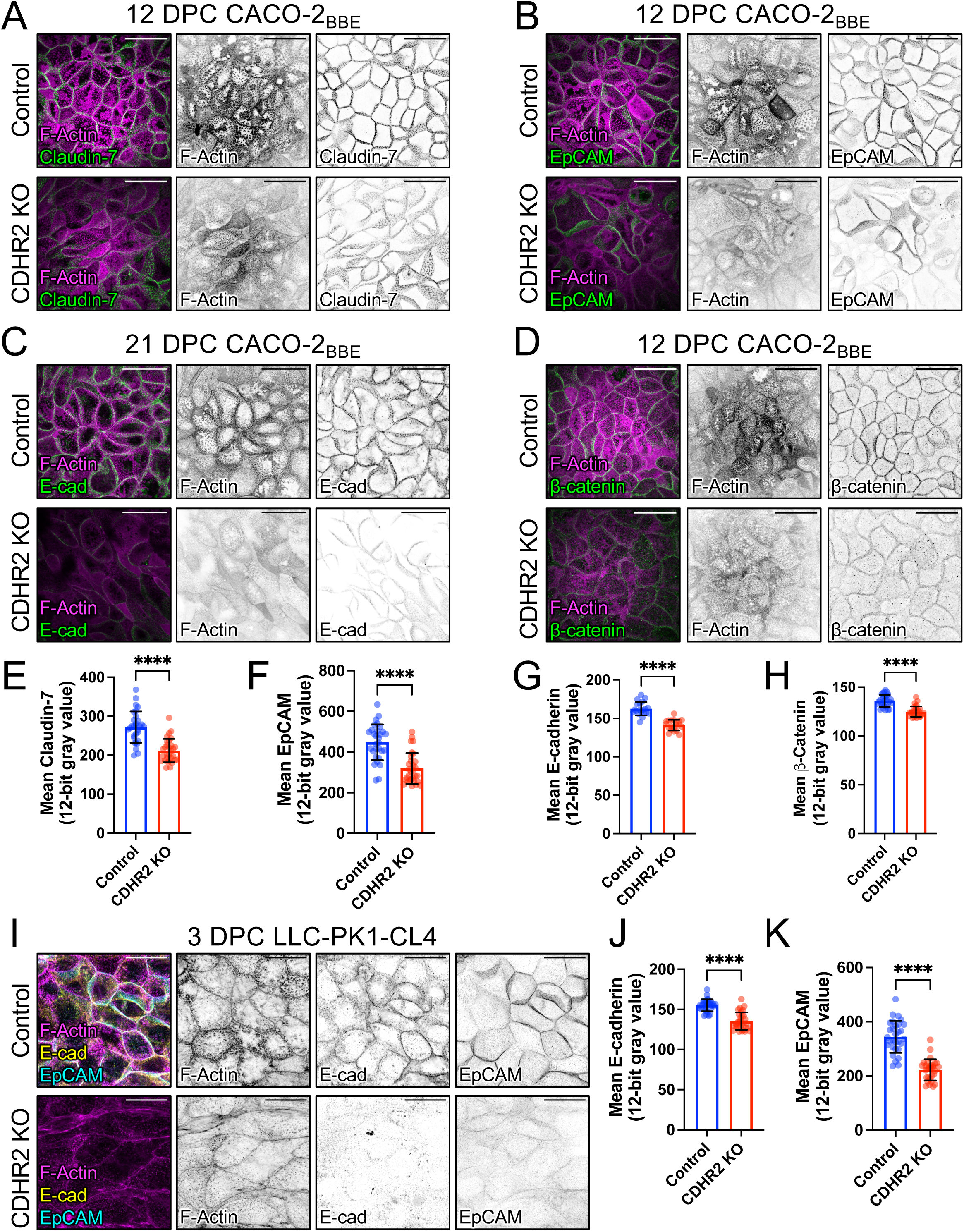
CDHR2 KO cells have decreased junction protein signal. (A) 12 DPC Control and CDHR2 KO CACO-2_BBE_ cells stained for F-actin (magenta) and Claudin-7 (green), and (B) F-actin (magenta) and EpCAM (green). (C) 21 DPC Control and CDHR2 KO CACO-2_BBE_ cells stained for F-actin (magenta) and E-cadherin (green). (D) 12 DPC Control and CDHR2 KO CACO-2_BBE_ cells stained for F-actin (magenta) and β-catenin (green). (E) Mean Claudin-7 intensities. (F) Mean EpCAM intensities. (G) Mean E-cadherin intensities. (H) Mean β-catenin intensities. (I) 3 DPC Control and CDHR2 KO CL4 cells stained for F-actin (magenta), E-cadherin (yellow), and EpCAM (cyan). (J) Mean E-cadherin and (K) EpCAM intensities. Intensity measurements for each junctional marker were taken from n = 30 60X fields per condition. Unpaired t-tests; ****p<0.005. Error bars represent mean ± SD. Scale bars: 40 µm (A-D) and 20 µm (I).

Our previous studies of the intestine-specific CDHR2 KO mouse focused on characterizing abnormalities in the apical brush border and revealed defects in microvillus structure and organization, as well as reduced IMAC enrichment at microvillar tips and decreased levels of apical solute transporters (Pinette et al., 2019). Based on our current findings, we revisited the CDHR2 KO mouse model to determine if defects in junctional composition also manifest *in vivo*. Indeed, staining and confocal imaging of paraffin sections from CDHR2 KO mouse small intestine revealed a marked loss of multiple TJ and AJ components, including ZO-1 (**Figure S3**), E-cadherin, EpCAM, claudin-7 and NM2C (**Figure 3**), and lower levels of the microvillar actin bundler, villin (**Figure S3D**) as noted in our previous report (Pinette et al., 2019). Inspection of the full length of the small intestine revealed that these phenotypes were most prominent at the duodenal end of the intestinal tract. In combination with our observations on CDHR2 KO CL4 and CACO-2_BBE_ models, we conclude that the molecular composition of canonical junctional complexes is disrupted in epithelial cells lacking CDHR2.

**Figure 3.**
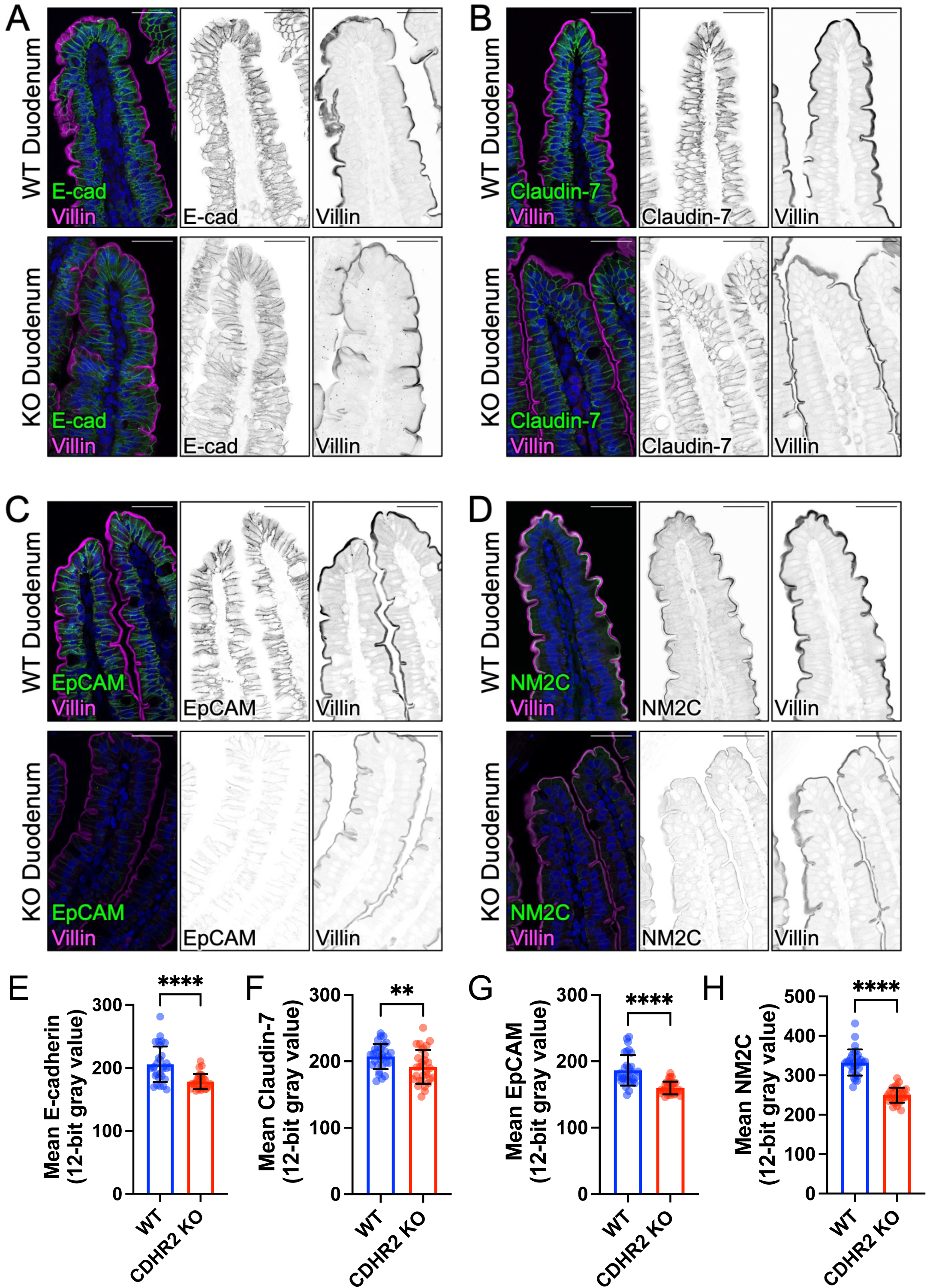
CDHR2 KO mouse duodenum exhibits decreased junctional protein signal. Wildtype and CDHR2 KO duodenum sections stained for villin (magenta) and the following markers in green: (A) E-cadherin, (B) Claudin-7, (C) EpCAM, (D) NM2C. Mean intensity measurements for WT vs. CDHR2 KO mouse for (E) E-cadherin, (F) Claudin-7, (G) EpCAM, (H) NM2C. Nuclei are marked by DRAQ5 (blue). Intensity measurements for each junctional marker were taken from n = 30 40X fields per condition, 2 WT and 2 CDHR2 KO littermates. Unpaired t-tests; **p = 0.0094, ****p<0.005. Error bars represent mean ± SD. Scale bars: 40 μm (A-D).

### CDHR2 KO cells exhibit a loss of junctional integrity and impaired wound healing

Based on the defects in junctional composition revealed by our quantitative immunostaining analysis, we next sought to determine if junctional integrity and function were impacted by CDHR2 KO. Toward this end, we turned to measurements of trans-epithelial electrical resistance (TEER), an established approach for quantitatively characterizing TJ formation and barrier integrity (Srinivasan et al., 2015). Control and CDHR2 KO CACO-2_BBE_ cells (two independently derived KO clones) were seeded on semi-permeable Transwell filters and TEER was measured every two days starting one day post-plating (**Figure 4A**). Measurements of TEER over the course of 22 days of differentiation revealed that, despite similar starting values, CDHR2 KO cells plateaued at a lower resistance compared to controls, with 22 DPC averages of 152.1 ± 12.2 Ω·cm^2^ vs. 342.6 ± 20.8 Ω·cm^2^, respectively (**Figure 4A-B**). Moreover, whereas the control cell TEER plateau fell within the range of resistance values reported in the literature for CACO-2 cells, the lower values measured for CDHR2 KO cultures indicate significant defects in TJ integrity (Narai et al., 1997; Srinivasan et al., 2015).

**Figure 4.**
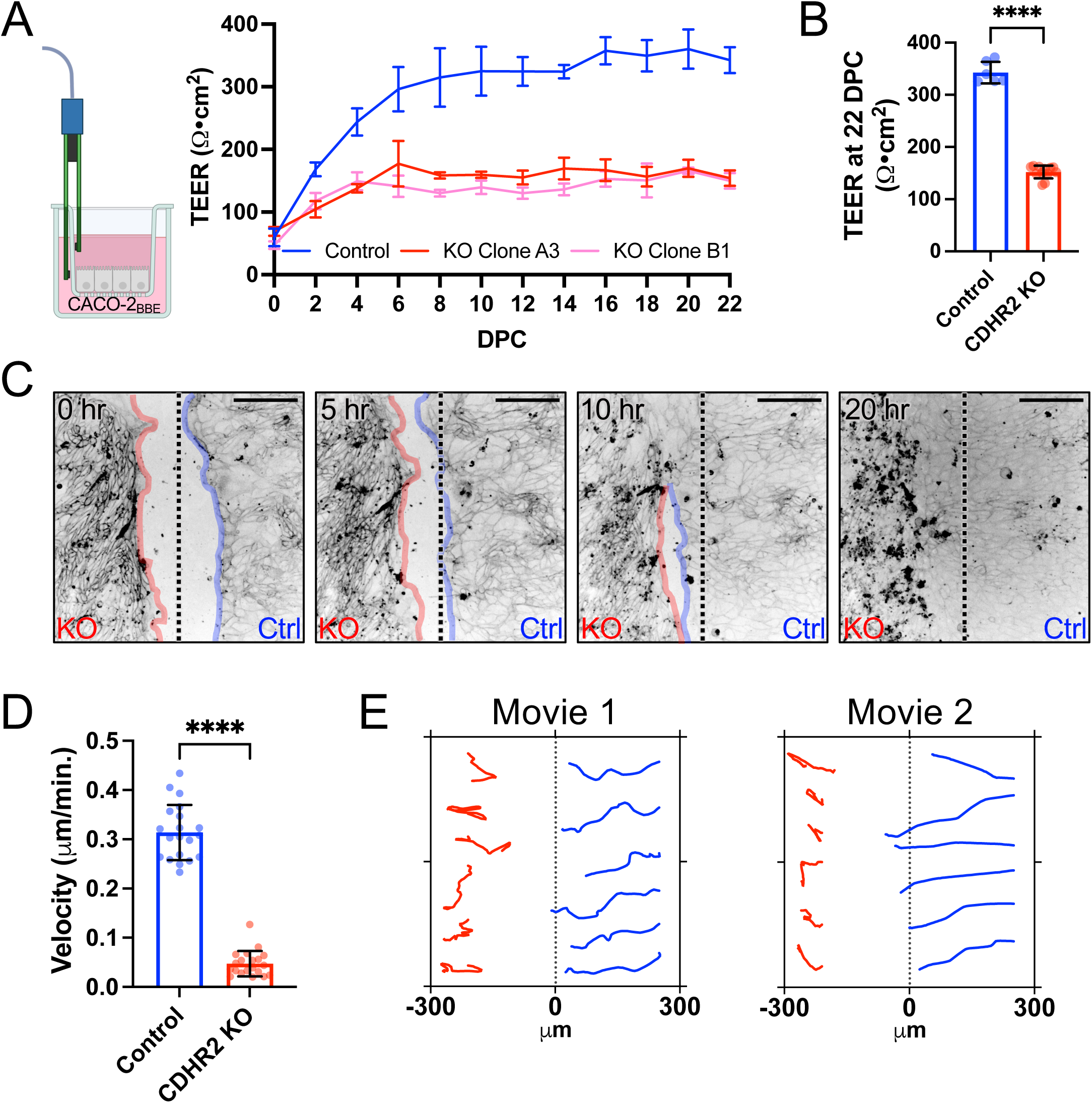
CDHR2 KO cells have impaired wound healing and decreased TEER. (A) CACO cells were seeded on Transwell inserts and transepithelial electrical resistance (TEER) measurements were measured every other day post-seeding to 22 DPC. (B) Mean TEER values from n = 6 Control and n = 12 CDHR2 KO (6 per KO clone) Transwells at 22 DPC: 342.6 ± 20.8 Ω·cm^2^ (Control) vs 152.1 ± 12.2 Ω·cm^2^ (KO clones). (C) Mixed Ibidi chamber cell migration assay with CDHR2 KO (left) and Control CL4 cells (right) +CellBrite650 membrane marker imaged over 20 hours. (D) Velocities of n = 20 Control and KO CL4 cell traces from the movie in (C) with mean velocities of 0.31 ± 0.06 μm/min (Control) and 0.05 ± 0.03 μm/min (KO); n = 20 lines drawn from the leading edge (see methods). (E) Individual traces of tracked cells from two replicate migration movies (Movie 1 shown in C). Unpaired t-test; ****p<0.005. Error bars represent mean ± SD. Scale bars: 200 µm (C).

The composition and integrity of cell junctions play key roles in defining monolayer-scale behaviors such as collective migration (De Pascalis and Etienne-Manneville, 2017; Gupta and Yap, 2021). To examine how loss of CDHR2 might impact this behavior, we seeded control and CDHR2 KO CL4 cells in chambers separated by a removeable barrier, which formed a wound-like gap on the surface once it was removed and allowed us to simultaneously observe the dynamics of both lines in the same imaging field. CL4 cell monolayers were labeled with a plasma membrane dye (CellBrite Steady 650), which enabled visualization of individual cell behaviors during attempted wound closure. Timelapse studies using this approach revealed that control CL4 cells exhibited well-ordered, collective migration towards the gap as expected (**Figure 4C-D and Movie S1**). In contrast, CDHR2 KO cells demonstrated almost no collective migration in the same timeframe, and velocity analysis confirmed a significant reduction in advance of the leading edge for KO cells relative to controls (0.05 ± 0.03 μm/min vs. 0.31 ± 0.06 μm/min, respectively (**Figure 4C-D**). Close examination of individual CDHR2 KO cells did show, however, that cells were still highly motile within the plane of the monolayer, although they demonstrated uncoordinated trajectories that rarely approached the gap (**Figure 4E**). Together, these functional assays indicate that loss of junctional complex components in CDHR2 KO cells leads to defects in regulated monolayer permeability and collective migration, two epithelial behaviors that are essential for physiological homeostasis.

### A role for intermicrovillar adhesion in stabilizing canonical junctional complexes

Given that transjunctional IMACs localize immediately above (i.e. apical to) TJs and AJs, we wondered if a loss of IMAC adhesion would impact the assembly and function of these canonical complexes. To test this idea, we characterized CDHR2 KO phenotypes in a newly developed CACO-2_BBE_ cell line and in a recently described CL4 line (Cencer et al., 2023). In both cell lines, CDHR2 KO led to defects in microvilli clustering, as expected (Crawley et al., 2014; Pinette et al., 2019). However, CDHR2 KO cells also presented with striking perturbations in cell and junction morphology. In the CL4 background, we observed significantly elongated cell profiles in confluent monolayers, a phenotype similar to previous reports in MDCK models lacking multiple junctional components (Choi et al., 2016). In the CACO-2_BBE_ background, CDHR2 KO led to a slightly different phenotype, although one still indicative of defects in junctional integrity. In this case, loss of CDHR2 led to a decrease in the straightness of individual junctional segments. One interpretation of this result is that ruffled (non-linear) junctions indicate a loss of mechanical tension, an idea consistent with the lower levels of NM2C that we observed at CDHR2 KO CACO-2_BBE_ junctions. Our staining experiments and quantitative imaging also revealed that CDHR2 KO in both cell line backgrounds led to a loss of core components from TJs and AJs; we noted similar perturbations in intestinal tissues from CDHR2 KO mice. These striking changes in NM2 enrichment and junctional composition could be linked, as previous studies revealed a complex interplay between normal levels of NM2-dependent tension and proper accumulation of F-actin and other core components of cell-cell contacts. For example, E-cadherin cell-cell contacts are strengthened under tension because of extracellular domain conformational changes that lead to catch bonding (Rakshit et al., 2012). Other studies showed that tension also stabilizes the E-cadherin/β-catenin binding and interactions with F-actin (Buckley et al., 2014). At this point, however, our data do not allow us to determine if the perturbations in junctional composition induced by CDHR2 KO are up or downstream of loss of NM2 and reduced junctional tension.

The changes in junctional morphology and composition induced by loss of CDHR2 were significant enough to disrupt the function of cell-cell contacts (**Figure 5**). Indeed, CDHR2 KO CACO-2_BBE_ cells exhibited a significant reduction in transepithelial electrical resistance, indicating that TJ integrity is compromised in the absence of normal IMAC function (Srinivasan et al., 2015; Yuan et al., 2020). Furthermore, wound healing assays showed that coordinated epithelial sheet motility was severely impaired in CDHR2 KO CL4 cell lines. These observations are consistent with recent work from others, which established that normal levels of ZO-1 are required for coherent epithelial migration (Skamrahl et al., 2021).

**Figure 5.**
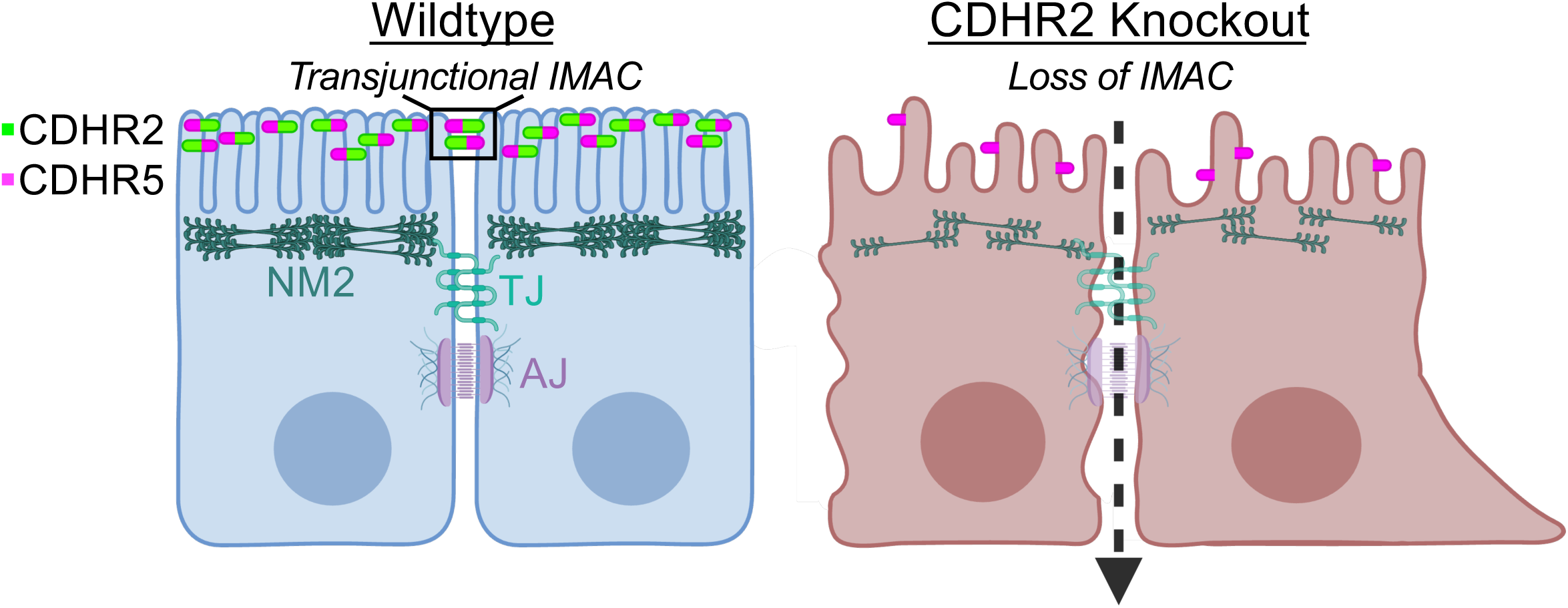
Loss of CDHR2-dependent intermicrovillar adhesion impairs basolateral junctions. Features of the CDHR2 KO phenotype include decreased microvillar clustering, aberrant cell morphology, reduced junctional tension, and a loss of key junctional proteins. These defects also give rise to increased junctional permeability and impaired collective cell migration during wound healing.

The findings we present here are consistent with some of the earliest literature on CDHR2 (initially referred to as ‘protocadherin LKC’), which proposed a tumor suppressor role for this factor (Okazaki et al., 2002; Ose et al., 2009). Early work showed that CDHR2 is downregulated in a range of colon and liver cancer cell lines (Okazaki et al., 2002). Later investigations reported that reintroduction of CDHR2 into HTC116 colon carcinoma cells drove dramatic changes in cellular phenotype, including recruitment of β-catenin to cell-cell contacts, accelerated wound healing, and suppression of tumor growth in mice (Ose et al., 2009). These findings led the authors to propose that β-catenin signaling downstream of CDHR2 drives contact inhibition, and loss of this activity promotes tumor formation in epithelial tissues. Consistent with those ideas, we found that CDHR2 KO in CACO-2_BBE_ demonstrated lower β-catenin levels at cell-cell contacts, a phenotype that has long been associated with aberrant epithelial cell motility and a loss of coordinated migration (Aman and Piotrowski, 2008).

Our group first discovered that CDHR2 participates in intermicrovillar adhesion on the apical surface of enterocytes in 2014 (Crawley et al., 2014). At that time, we were perplexed by the early studies invoking a tumor suppressor role in intestinal tissues, because they implied that CDHR2 was directly involved in β-catenin stabilization at junctional complexes (Okazaki et al., 2002; Ose et al., 2009). That model was difficult to reconcile with super-resolution images showing that CDHR2 instead targets specifically to the distal tips of brush border microvilli (Crawley et al., 2014). A resolution to this conundrum emerged with the discovery that CDHR2 forms IMACs between microvilli that extend from neighboring cells (Cencer et al., 2023). Based on the data we report here, we propose that these transjunctional IMACs form the most apical layer of physical contact between neighboring epithelial cells and promote the integrity of more basolateral adhesion complexes, including TJs and AJs. Future studies seeking to build on these results might focus on the role of transjunctional IMACs in response to challenges that arise in intestinal pathologies, such as Crohn’s disease (VanDussen et al., 2018).

## ACKNOWLEDGMENTS

The authors thank all members of the Tyska laboratory for their constructive feedback. We thank the Vanderbilt Genome Editing Resource (VGER) supported by the Cancer Center Support Grant P30 CA68485 for their expertise and assistance in generating the CDHR2 knockout mouse. We also acknowledge the Translational Pathology Shared Resource supported by NCI/NIH Cancer Center Support Grant P30 CA068485 for their assistance in paraffin embedded tissue sectioning. FACS was performed in the VUMC Flow Cytometry Shared Resource supported by the Vanderbilt Ingram Cancer Center P30 CA68485, and the Vanderbilt Digestive Disease Research Center supported by P30 DK058404. This study was also supported by the NIH grants R01 DK095811 (M.J.T.), R01 DK125546 (M.J.T.), and R01 DK111949 (M.J.T.), and the Training Program in Developmental Biology T32 HD007502 (C.V.W).

## SUPPLEMENTAL MOVIE LEGENDS

**Movie S1, related to Fig. 4**. Collective cell migration and wound healing is impaired in the absence of CDHR2. CDHR2 KO (left) and Control (right) CL4 cells after Ibidi chamber removal marked with the membrane dye CellBrite Steady 650 imaged over ∼22 hours at 25-minute intervals. Scale bar represents 200 μm.

## MATERIALS AND METHODS

### Animal models

The CDHR2 knockout (KO) mouse model was previously created and characterized by our laboratory (Pinette et al., 2019). Animal experiments were carried out in accordance with Vanderbilt University Medical Center Institutional Animal Care and Use Committee guidelines under IACUC Protocol M1600206-02.

### Cell culture models

LLC-PK1-CL4 porcine kidney proximal tubule cells were grown in 1X high glucose DMEM/2mM L-glutamine (Corning #10-013-CV) with an added 1% L-glutamine (Corning # 25-005-CI) and 10% fetal bovine serum (FBS) (R&D Systems). CACO-2_BBE_ human colonic adenocarcinoma cells were grown in the same medium, with 20% FBS. Cells were incubated at 37°C and 5% CO_2_. Regular mycoplasma testing was performed using the MycoStrip Mycoplasma Dectection Kit (InvivoGen #rep-mys-50).

### CRISPR Knockout

The CDHR2 CL4 KO cell line was created and validated in our previous study (Cencer et al., 2023). The CDHR2 CACO-2_BBE_ KO cell line was generated and validated for the present study also as previously described, using the Lenti-CRISPR v2 system, with gRNA’s designed to target exons 3 and 4 of human CDHR2 genomic sequence: [Exon 3 FWD CACCGTAGGAACTTCGGGGCCACGT; Exon 3 REV AAACACGTGGCCCCGAAGTTCCTAC; Exon 4 FWD CACCGTCTTCCGCTACCAACCAGA; Exon 4 REV AAACTCTGGTTGGTAGCGGAAGAC] PCR/sequencing primers targeted regions surrounding Exon 3 or Exon 4, within which Cas9 was predicted to cut: [Exon 3 Fwd CAGCTATGGCTGTCCTGCTTC; Exon 3 Rev CAGTCAGAGACTGAAAGCGATGG; Exon 4 Fwd CTCCAAAGCTCTAGTCTGCACC; Exon 4 Rev CTCACCTGGATGTAGGGGTCG]

### Rescue CL4 Cell Line Generation

A validated CDHR2 KO CL4 cell clonal population was transfected with pHALO-N3-CDHR2 using FuGENE 6 (Promega #E2691) at a FuGENE:DNA (μL:μg) ratio of 3:1 following the reagent protocol in a T25 cell culture flask. The next day, cells were split to a T75 flask plus 1mg/mL G418 sulfate for antibiotic selection. Cells were maintained in culture under G418 selection to create a stable cell line.

### Fluorescence-activated cell sorting (FACS)

CDHR2 KO/+HALO-CDHR2 rescue CL4 cells were incubated with 50nM Janelia Fluor 635 dye for 1 hour at 37°C to label cells, trypsinized, and sorted for mid-high expression as previously described by Vanderbilt University Medical Center’s Flow Cytometry Shared Resource on a 5-Laser FACS Aria III system with a 100 µm sized nozzle (Cencer et al., 2023). Sorted cells were maintained under 1mg/mL G418 selection to maintain stable expression.

### Swiss roll and paraffin embedded tissue preparation

The entire mouse small intestinal tube was excised and flushed with cold 1X PBS. Tissue was fixed in the tube using the hemostat technique described above with room temperature 2% PFA for 15 minutes. After removing the hemostats, the intestinal tube was slid onto a metal cannula and cut lengthwise, down the entire length, with scissors. The flayed tissue was then rolled out, villi side up, onto a strip of parafilm. A hemostat was clamped onto the proximal end (duodenum) of the intestine and rolled, with the duodenum at the center of the roll. A 21g needle was stuck through the roll and the hemostat was removed. The roll was then submerged in 10% neutral buffered formalin at room temperature for 48 hours. After fixation, the needle was removed and the roll was cut in half, creating two thinner rolls, and each half was placed into a large tissue cassette and submerged back into the formalin. Cassettes were submitted to the Vanderbilt University Translational Pathology Shared Resource, embedded in paraffin wax, and sliced onto glass slides, at 10 µm thickness. Slides were stored at room temperature until staining.

### Paraffin embedded tissue staining

Using a Tissue-TekII manual slide staining set, slides were deparaffinized in Histo-Clear II (National Diagnostics) 2 times, 3 minutes each. Tissue was then rehydrated in a descending ethanol series [100%, 100%, 95%, 90%, 70%, 50%] 5 minutes each followed by washing in PBS 3 times, 3 minutes each. Slides were incubated in antigen retrieval buffer [10mM Tris, 0.5mM EGTA, pH 9.0] in coplin jars for 1 hour using a rice cooker and then cooled to room temperature. Slides were washed 3 times, 3 minutes each in PBS and then blocked in 10% NGS for 1 hour at room temperature. Primary antibody (diluted in 1% NGS) was added overnight at 4°C. The next day, slides were washed 3 times, 5 minutes each in PBS and secondary antibody (diluted in 1% NGS) was added for 1 hour at room temperature in the dark. Slides were then washed 3 times, 5 minutes each in PBS followed by dehydration with an ascending ethanol series [50%, 70%, 90%, 95%, 100%, 100%] 5 minutes each. A coverslip was mounted with ProLong Gold. The following antibodies and dilutions were used for paraffin section staining: anti-ZO-1 (rabbit, Invitrogen #61-7300), 1:50; anti-Villin (mouse, Santa Cruz #SC-66022), 1:50; or anti-Villin (rabbit, Santa Cruz # SC-28283) 1:50; anti-E-cadherin (mouse, BD Biosciences #610182), 1:100; anti-EpCAM (rabbit, Invitrogen #PA5-19832), 1:100; anti-MYH14/NM2C (rabbit, Proteintech #20716-1-AP), 1:100; anti-Claudin-7 (rabbit, Invitrogen #34-9100), 1:100; Alexa Fluor F(ab’)2 fragment goat anti-rabbit 488 (Invitrogen #A11070), 1:1000; Alexa Fluor goat anti-mouse 568 (Invitrogen #A11019), 1:1000; Alexa Fluor F(ab’)2 fragment goat anti-rabbit 568 (Invitrogen #A21069), 1:1000; Alexa Fluor F(ab’)2 fragment goat anti-mouse 488 (Invitrogen #A11017), 1:1000. The secondary antibodies were spun down for 10 minutes at 4°C and 21 x g prior to using. DRAQ5 was used to label nuclei (Molecular Probes #62251); 1:500.

### Fixed cell immunofluorescence

CL4 and CACO-2_BBE_ cells were grown to *n* days post-confluent (DPC) on acid-washed 22x22 mm #1.5H coverslips (Globe Scientific) in a 6-well plate to a polarization time point of 3 DPC and 12 or 21 DPC, respectively. Cells were fixed and stained as previously described (Cencer et al., 2023). The following antibodies and dilutions were used for cell staining: anti-PCLKC (CDHR2) (mouse, Abnova #H00054825-M01), 1:25; anti-CDHR5 (Rabbit, Sigma #HPA009173), 1:250; anti-ZO-1 clone R40.76 in CL4 (rat, EMD Millipore Sigma #MABT11), 1:100; anti-ZO-1 in CACO-2_BBE_ (rabbit, Invitrogen #61-7300), 1:50; anti-E-cadherin (mouse, BD Biosciences #610182), 1:100; anti-EpCAM (rabbit, Invitrogen #PA5-19832), 1:100; anti-MYH14/NM2C (rabbit, Proteintech #20716-1-AP), 1:100; anti-Claudin-7 (rabbit, Invitrogen #34-9100), 1:100; anti-Beta-Catenin (rabbit, Invitrogen #71-2700), 1:100; Alexa Fluor F(ab’)2 fragment goat anti-rabbit 488 (Invitrogen #A11070), 1:1000; Alexa Fluor goat anti-mouse 568 (Invitrogen #A11019), 1:1000; Alexa Fluor F(ab’)2 fragment goat anti-mouse 488 (Invitrogen #A11017) and goat anti-rabbit 568 (Invitrogen #A21069), 1:1000; Alexa Fluor goat anti-rat 647 (Invitrogen #A21247), 1:200; and Alexa Fluor Plus 405 Phalloidin (Invitrogen # A30104) or Alexa Fluor 647 Phalloidin (Invitrogen # A22287), 1:200 for actin staining. The secondary antibodies, not including phalloidin, were spun down for 10 minutes at 4°C and 21 x g prior to using.

### Fixed sample microscopy

Laser scanning confocal microscopy was performed on a Nikon A1 microscope equipped with 488 nm, 568 nm, and 647 nm LASERs using an Apo TIRF 100x/1.49 NA or Apo TIRF 60x/1.49 NA TIRF oil immersion objective. Spinning disk confocal microscopy was used for CRISPR CDHR2 KO cell and CDHR2 KO mouse tissue imaging and intensity analysis using a Nikon Ti2 inverted light microscope with a Yokogawa CSU-W1 spinning disk head, a Photometrics Prime 95B sCMOS camera, and four excitation LASERs (488, 568, 647, and 405 nm) and a 60X/1.49 NA TIRF oil immersion objective.

### Wound Healing Assays

<u>CL4 cells</u> were seeded at a total of 30,000 cells per well in an Ibidi 2-chamber insert (Ibidi #80209) adhered to a 35 mm plasma cleaned glass-bottom dish (CellVis #D35-20-1.5-N). Cells were grown for 2 days or until they had just reached the edge of the chamber. The chamber media was aspirated, and the insert was carefully removed with forceps. Cells were gently rinsed twice with warm DPBS and fresh media was added to the dish. Cells were left in the incubator overnight, ∼12 hours, to recover. Before imaging, cells were rinsed and 1:1000 CellBrite Steady 650 membrane dye and 1:1000 kit enhancer (Biotium #30108) was added in Fluorobrite DMEM +10% FBS for 30 minutes at 37℃.

Cells were then imaged using a 10X Plan Apo 0.45 NA objective on a Yokogawa CSU-X1 spinning disk confocal microscope with a 647 nm excitation LASER for ∼24 hours, or until 100% wound closure.

<u>CACO-2_BBE_ cells</u> were seeded at a total of 50,000 cells per well of a 24 well plate (Thermo Scientific #142475) and allowed to grow until they just reached 100% monolayer coverage. Each monolayer was scratched in a single line spanning the well, top to bottom, with a P10 pipette tip and wound closure was imaged using a BioTek Cytation 5 plate reader in 1-hour intervals over a total of 20 hours.

### Transepithelial Electrical Resistance

Transwells (Greiner Bio-one #662641) were primed with 100 μL of cell culture media in the top chamber and 600 μL in the bottom chamber and left at 37°C for 15 minutes prior to cell seeding. Control and CDHR2 KO clone CACO-2_BBE_ cells were counted, and 30,000 cells were seeded in each transwell with a total volume of 100 μL. A “blank” transwell was also maintained with media alone for background TEER measurements. 24 hours post-seeding, the first TEER measurements were taken for the “0 DPC” time point. TEER was measured in ohms (Ω) using an EVOM3 epithelial voltohmmeter device (World Precision Instruments) equipped with a calibrated electrode (World Precision Instruments #STX2-PLUS). Prior to each TEER measurement, existing media was exchanged for fresh media, and cells were incubated for 4 hours to ensure equal volume and equilibration in each well. The raw blank TEER value was subtracted from each monolayer TEER value and then multiplied by 0.33 cm^2^, the area of the transwell, to obtain the reported TEER value in Ω·cm^2^.

## QUANTIFICATION AND STATISTICAL ANALYSIS

### CDHR2 KO cell intensity measurements

A total of 45 60X fields were used for each condition, control or KO, from stained coverslips. (For the KO, 15 60X fields were taken for each sequenced clone, for a compiled total of 45 KO fields.) Raw images were maximally projected in Z in FIJI and mean intensity of the entire field was measured for the CDHR2 and CDHR5 channels.

The measured intensities were plotted in Prism and statistics were quantified with an unpaired t-test.

### CL4 cell shape measurements

60X fields of 3DPC Control and CDHR2 KO CL4 cells stained for ZO-1 were analyzed in Elements using a custom General Analsysis 3 (GA3) pipeline. This involved first maximally projecting each image in Z followed by tight border segmentation. The binary was then inverted to highlight individual cells in the field and any “partial” cells touching the borders were eliminated from the selection. Cell area (μm^2^) and elongation were measured from the remaining binaries. Elongation is quantified by the software as a ratio of maximum feret length over minimum feret length. A value of greater than 1 indicates that the cell is stretched in one of its axes.

### CACO cell junction straightness measurements

Three randomly selected 60X fields were taken from each of three experimental staining replicates and maximally projected in FIJI. The ZO-1 channel was isolated and junctional segments (between two vertices) were cropped using the rectangle selection tool. The cropped segment was then binarized and dilated to segment the ZO-1 signal marking the cell junction. Next, the binary was skeletonized and the 2D skeleton was analyzed. Branch length and Euclidian distance were exported from the measurements list to Excel. Junction straightness was calculated as the ratio of Euclidean distance over the branch length, with 1 being the most straight, as previously described in the literature (Sumi et al., 2018). A total of 62 straightness ratios were calculated and plotted in Prism as a column data table and graphed as a violin plot. An unpaired t-test was performed to statistically compare the mean straightness of junctions in Control vs KO cells.

### Wound healing velocity measurements

Cell migration movies were maximally projected in FIJI and divided in half with a duplicated ROI. For the Control side, 20 randomly placed horizontal lines were drawn along the entire vertical axis of the leading edge of the cell monolayer to the opposite end of the ROI (the wound “midline”). The length, in μm, was recorded in Excel along with the time, minutes, it took to reach the midline and velocity was calculated from μm/min. For the KO side, which exhibited far less movement towards the wound midline, 20 randomly placed horizontal lines were drawn from the monolayer edge to the furthest point of migration. Both length of the line and time in the movie were recorded and velocity was calculated from μm/min. Each velocity measurement was plotted in FIJI in a column data sheet. An unpaired t-test was performed to test significance of the change in mean velocities between Control and KO cells.

### Tissue intensity measurements

For the quantification of staining levels, 15 40X 1.39NA images (Nikon W1 spinning disk) per mouse (two control, two wildtype littermates) were analyzed for mean intensity using a Nikon Elements GA3 analysis pipeline. Broadly, each raw image was maximally projected in Z and the junctional marker channel was thresholded for intensity encompassing respective signal in control mice. If necessary, to encompass all junctional signal, the thresholded ROIs were dilated and/or cleaned. The same thresholding parameters were applied to CDHR2 KO mouse images to maintain comparability in the analysis. Mean intensity readouts were exported from Elements and analyzed using Prism. An unpaired t-test was used to compare mean junction intensity.

