## Supplemental information for "Loss of intermicrovillar adhesion impairs basolateral junctional complexes in transporting epithelia"

**Supplemental Figures**

**Figure S1, related to Fig. 1.** Generation and validation of CDHR2 KO CACO-$2_{BBE}$ cells.

**Figure S2, related to Fig. 2.** CDHR2 KO CL4 cells have reduced CDHR5 and ZO-1 that is rescued upon exogenous CDHR2 expression.

**Figure S3, related to Fig. 3** Tight junction protein ZO-1 and actin bundling protein villin are depleted in the CDHR2 KO mouse small intestine (duodenum).

**Supplemental Movies**

**Movie S1, related to Fig. 4** Collective cell migration and wound healing is impaired in the absence of CDHR2. CDHR2 KO (left) and Control (right) CL4 cells after Ibidi chamber removal marked with the membrane dye CellBrite Steady 650 imaged over ~22 hours at 25-minute intervals. Scale bar represents 200 µm.

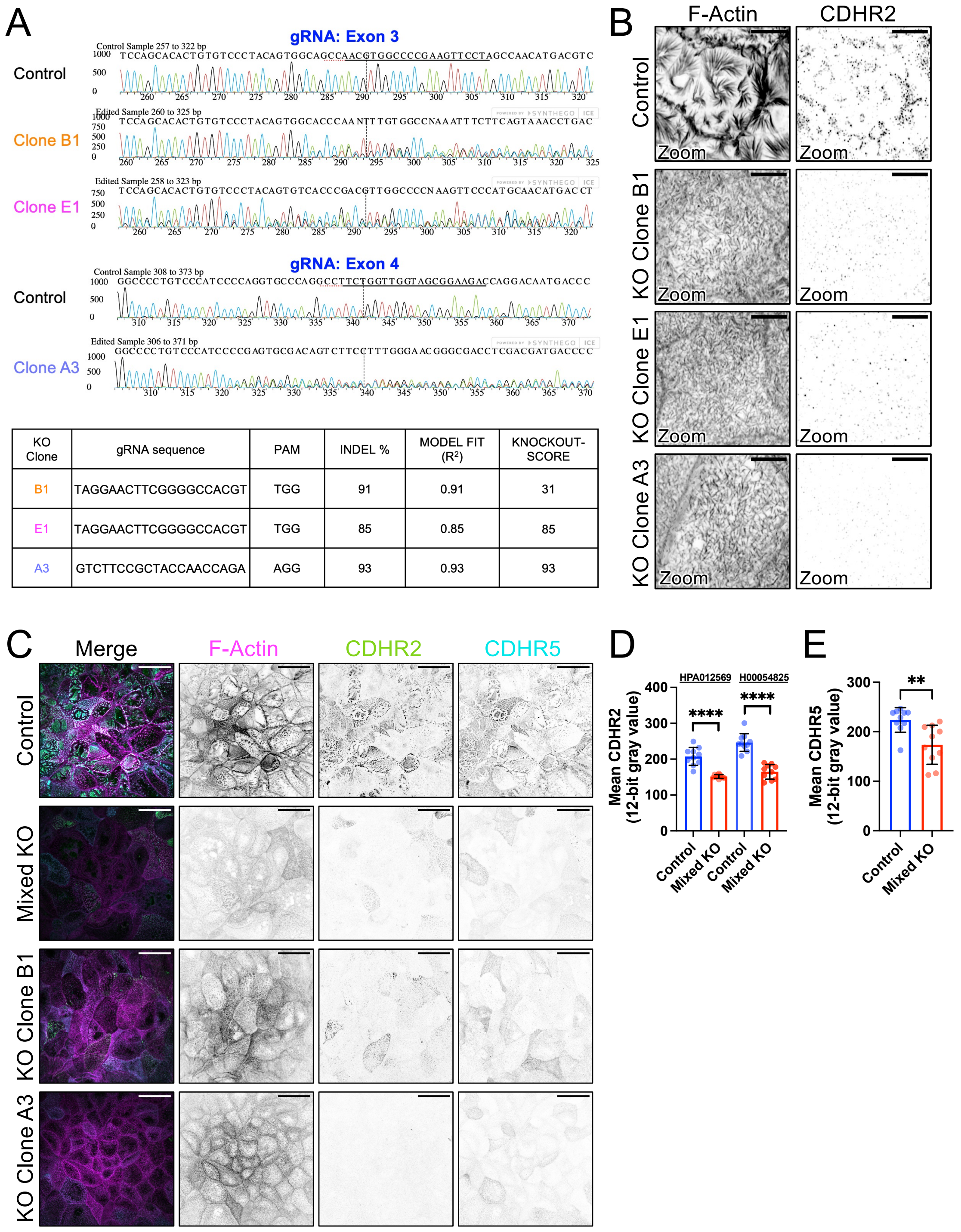


| KO Clone | gRNA sequence | PAM | INDEL % | MODEL FIT ($R^2$) | KNOCKOUT-SCORE |
|---|---|---|---|---|---|
| B1 | TAGGAACTTCGGGGCCACGT | TGG | 91 | 0.91 | 31 |
| E1 | TAGGAACTTCGGGGCCACGT | TGG | 85 | 0.85 | 85 |
| A3 | GTCTTCCGCTACCAACCAGA | AGG | 93 | 0.93 | 93 |

**Figure S1, related to Figure 1.** Generation and validation of CDHR2 KO CACO-$2_{BBE}$ cells. (A) Genomic DNA was extracted from selected KO clonal populations and PCR was used to generate two regions spanning exons 3 and 4 of CDHR2. Trace files of each clone and the control cells were analyzed with the Synthego Inference of CRISPR Edits (ICE) tool (Conant et al., 2022). (B) Laser scanning confocal MaxIP images of Control, CDHR2 KO Clone "B1", "E1", and "A3" stained for F-Actin and CDHR2 as labeled showing lack of microvillar clustering in the KO cells. (C) W1 spinning disk MaxIP images of Control, mixed KO (pre-clonal isolation), and KO clones stained for F-Actin (magenta), CDHR2 (green), and CDHR5 (cyan). (D) Mean CDHR2 intensities of Control vs. CDHR2 mixed KO using two different CDHR2 antibodies, as marked. (E) Mean CDHR5 intensities of Control vs. CDHR2 mixed KO. n = 10 60X fields per condition. Unpaired t-test; **** p ≤ 0.0001, **p = 0.0032. Error bars represent mean ± SD. Scale bars: 5 µm (B), 30 µm (C).

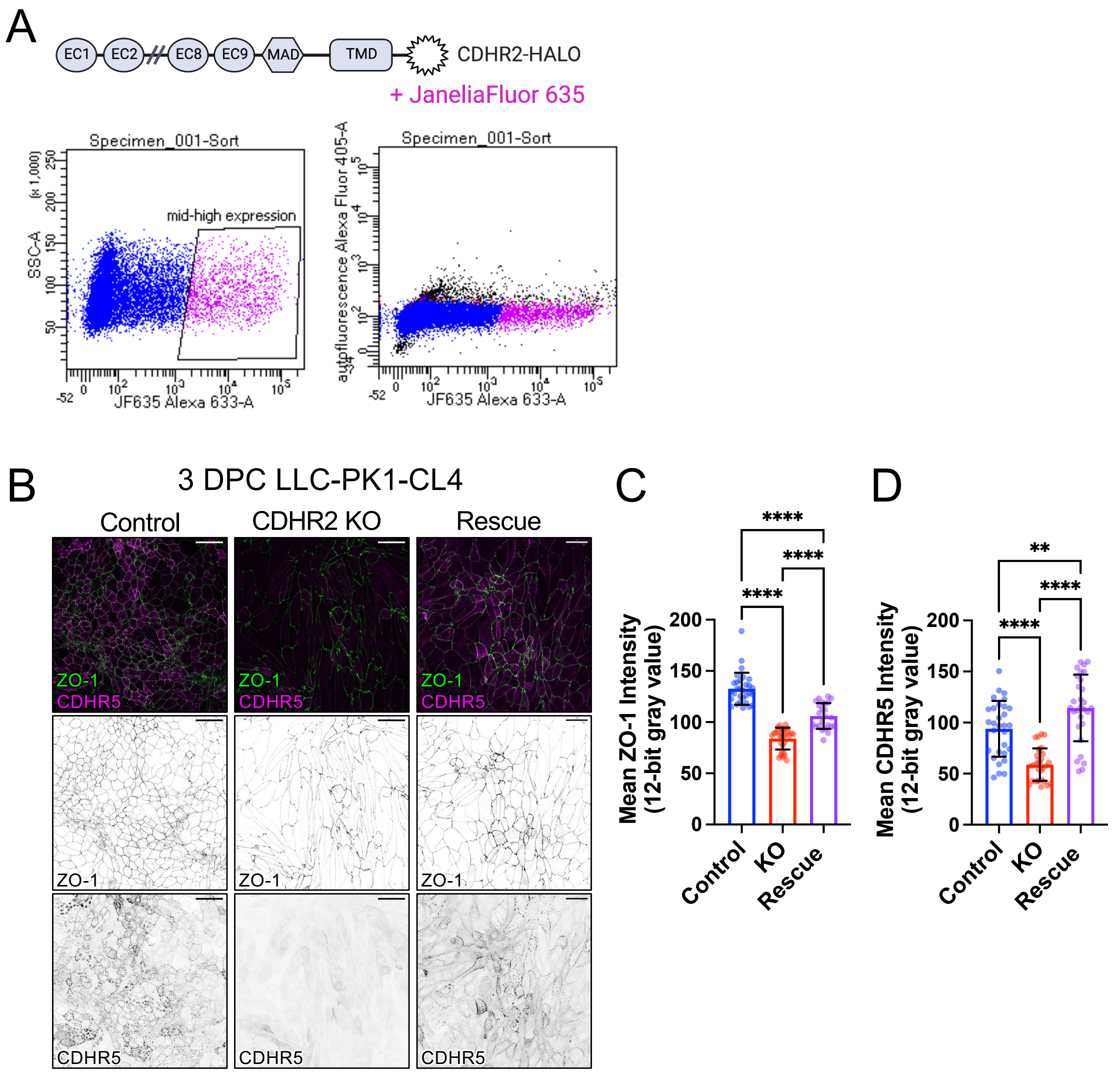


**Figure S2, related to Figure 2.** CDHR2 KO CL4 cells have reduced CDHR5 and ZO-1 that is rescued upon exogenous CDHR2 expression. (A) Diagram of the C- tagged CDHR2-HALO construct. “Rescue” cells were transfected, stably selected for CDHR2-HALO and isolated via FACS using the HALO ligand JF635. (B) Scanning laser confocal MaxIPs of 3 DPC Control, CDHR2 KO, and CDHR2-HALO rescue CL4 cells stained for ZO-1 (green) and CDHR5 (magenta). (C) Mean CDHR5 and (D) mean ZO-1 intensities for the three cell conditions. n = 30 imaged 40X fields per condition from 3 independent staining experiments. $^{**}p = 0.0095$, $^{****}p \leq 0.0001$ Ordinary one-way ANOVA with post-hoc multiple comparisons test. Error bars represent mean ± SD. Scale bars: 40 μm (B).

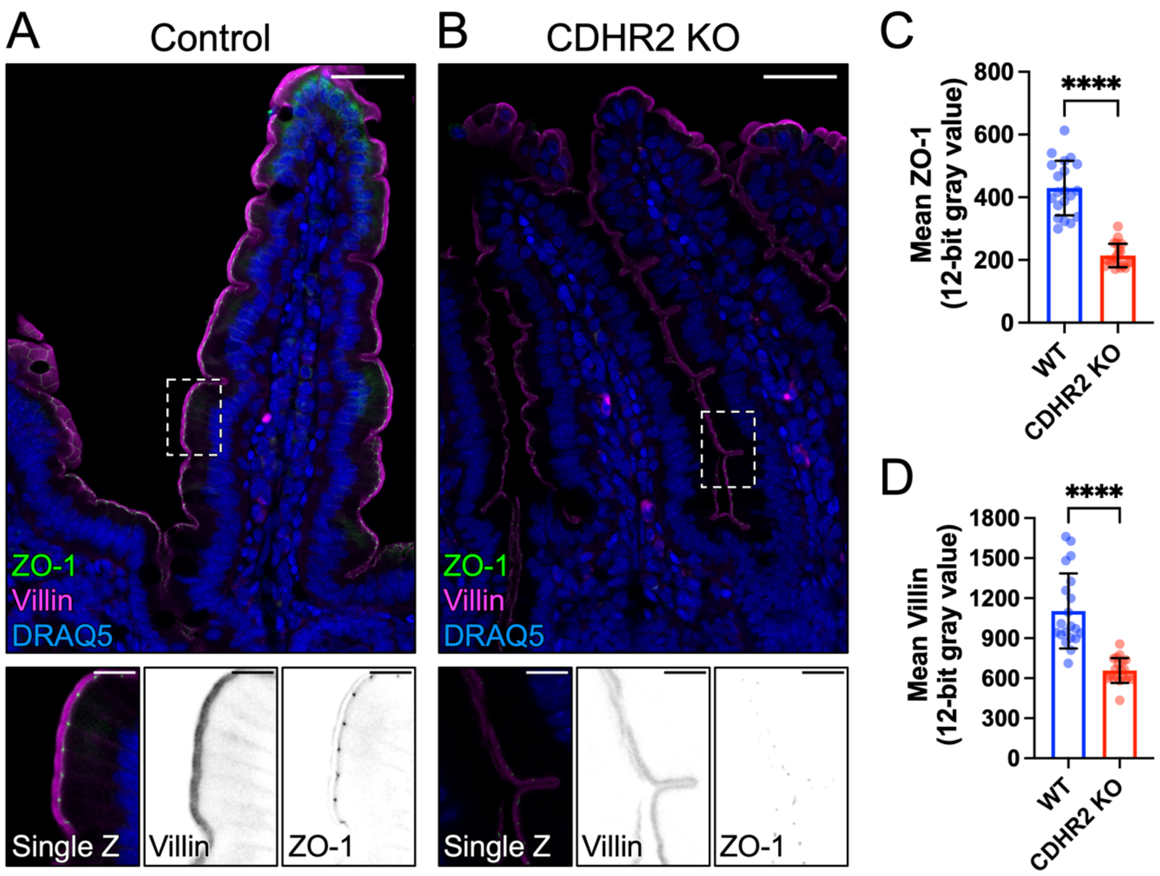


**Figure S3, related to Figure 3.** Tight junction protein ZO-1 and actin bundling protein villin are depleted in the CDHR2 KO mouse small intestine (duodenum). Laser scanning confocal MaxIPs of 10 μm paraffin sections stained for ZO-1 (green), villin (magenta), and DRAQ-5 (blue) from (A) Control (Cre-) and (B) CDRH2 KO (Cre+) mice. The dashed boxes represent the zoom area shown below as a single Z-slice with merged and inverted channel images. LUTs are matched/scaled to Control images. (C) Thresholded mean ZO-1 intensity measurements and (D) thresholded mean villin intensity measurements from n = 2 Control and n = 2 CDHR2 KO mice littermates, 20 measured villi per condition. **** $p \leq 0.0001$ unpaired t-test. Error bars represent mean ± SD. Scale bars: 40 μm (A-B), 10 μm (Zooms).
